# Tameness selection pressure affects gut virome diversity in mice

**DOI:** 10.1101/2024.08.26.609628

**Authors:** Bhim B. Biswa, Kazumichi Fujiwara, Atsushi Toyoda, Tsuyoshi Koide

## Abstract

The gut microbiome, a complex ecosystem comprising bacteria, viruses, archaea, fungi, and other microorganisms, plays a pivotal role in host health, immunity, and behaviour. Among its components, bacteriophages and viruses that infect bacteria significantly influence microbial community dynamics by affecting bacterial diversity, function, and evolution. Despite extensive research on bacterial components, the viral metagenome (virome) remains relatively unexplored. This study investigated the effect of selective breeding for tameness on the gut virome of wild heterogeneous stock (WHS) mice. WHS mice were selectively bred for active tameness, resulting in four groups: S1 and S2 (selected for tameness), and C1 and C2 (non-selected). In previous study, we observed an increased abundance of *Limosilactobacillus reuteri* in tame groups compared to non-selected groups, as well as a tameness-promoting effect of *L. reuteri* with long-term administration. Given the regulatory role of phages in bacterial populations, this study focused on analysing the gut virome using shotgun metagenome sequencing data. From the 84 samples, we generated 6078 non-redundant viral operational taxonomic units (vOTUs) and identified the hosts of 3,065 of these vOTUs. Significant differences in gut virome beta diversity were observed between the selectively bred and control groups, suggesting that tameness selection exerts distinct pressure on the virome. Additionally, phage-host interaction analysis revealed strong correlations between specific phages and their bacterial hosts, indicating a co-occurrence that influences host behaviour. Overall, this study provides novel insights into the role of the gut virome in shaping host behaviour and highlights the broader implications of microbial contributions to domestication and selective breeding outcomes.

## Introduction

The gut microbiome is a complex ecosystem comprising not only bacteria, but also viruses, archaea, fungi, and other microorganisms. Bacteriophages or viruses that infect bacteria play pivotal roles in shaping microbial communities by influencing their diversity, function, and evolution[1, 2]. Despite extensive research on the bacterial components of the gut microbiome, the virome, particularly the viral metagenome, remains relatively unexplored. Elucidating the gut virome is crucial because it can have a profound impact on host animal health, immunity, and behaviour through its interactions with bacterial hosts.

Domestication and selective breeding have long been known to alter animal behaviour and physiology, primarily through genetic modifications. The role of the gut microbiome, including its viral components, in these processes has gained increasing attention. Selective breeding, especially for traits such as tameness, involves sustained human intervention that may influence the composition and function of gut microbial communities, including the virome. Recent studies suggest that the gut microbiota can modulate host behaviour[3, 4], with potential contributions from gut-resident viruses[5, 6].

One of the most important parameters for animal domestication is tameness, which can be divided into two types: active (motivation to interact with humans) and passive (reluctant to avoid humans)[7]. Previously it was found that laboratory mice show passive tameness[8]. To develop a mouse line displaying active tameness and study domestication, wild heterogeneous stock (WHS) mice were established by crossbreeding eight different wild strains of mice (BFM/2Ms, PGN2/Ms, HMI/Ms, NJL/Ms, BLG2/Ms, KJR/Ms, CHD/Ms, and MSM/Ms) collected from different part of the world. Active tameness selection was applied to this line as a selection pressure over generations[8], resulting in four independent mouse groups: S1, S2, C1, and C2. S1 and S2 were subjected to selective breeding against active tameness, whereas C1 and C2 were subjected to random breeding without the selection pressure and served as controls[9]. Genomic loci under selection due to active tameness selection pressure were identified through selective mapping of these groups[10]. Subsequently, the focus shifted to the gut microbiome and its potential role in tameness behaviour. The abundance of *Limosilactobacillus reuteri* was significantly higher in the tame groups, and administration of *L. reuteri* to the control groups increased their tameness[11]. Given that phages usually influence bacterial populations[12, 13], the present study focused on the gut virome.

This study aimed to explore the influence of selective breeding on the gut virome of WHS mice by analysing viral metagenome-assembled genomes (MAGs) from shotgun metagenome sequence data[11]. We hypothesised that selective breeding for tameness not only affects the bacterial community but also alters the gut virome, potentially contributing to the observed behavioural changes[11, 14]. By characterising viral MAGs, we sought to uncover novel interactions between bacteriophages and their bacterial hosts that may play a role in the domestication of host animals. We conducted a comprehensive analysis of viral MAGs isolated from the gut microbiota of selectively bred and control WHS mice. Our approach included advanced bioinformatics techniques to identify and classify viral sequences, evaluate their diversity and abundance, and examine their potential impact on host and bacterial communities. This study provides new insights into the role of the gut virome in shaping host behaviour and offers a broader understanding of microbial contributions to domestication and selective breeding outcomes.

## Materials and Methods

### Experimental conditions

All experiments were conducted in strict accordance with the guidelines and protocols approved by the “Committee for Animal Care and Use” of the National Institute of Genetics (permit numbers R4-25 and R5-7). The mice used in our experiments were bred and maintained under specific pathogen-free (SPF) conditions in the animal facility at the NIG. In this study, 27^th^ generation WHS line was used.

### Selective Breeding of WHS Mice

WHS mice were generated by crossing eight wild strains (BFM/2Ms, PGN2/Ms, HMI/Ms, NJL/Ms, BLG2/Ms, KJR/Ms, CHD/Ms, and MSM/Ms) from different countries to enhance their genetic heterogeneity [10]. The founder stock, maintained with 16 breeding pairs, was split into two groups in the third generation: S1 and C1. S1 was selectively bred for greater active tameness. In the fifth generation, another selected group (S2) was split from C1, and another non-selected group (C2) was split from S1. Subsequently, S2 was selectively bred for higher levels of active tameness, whereas C2 was maintained as the non-selected group. For breeding, mice were selected based on the highest contact score in the active tameness test from five female and five male offspring in each pair. If multiple animals had the same contact score, the animal with the highest heading score in the same test was chosen.

### Tameness test and faeces collection

Tameness tests and faecal collection were performed as described previously [9, 11, 15].

### Metagenomic DNA isolation and sequencing

Metagenomic DNA was isolated as previously described [11]. Briefly, metagenomic DNA was isolated from faeces using lysis buffer (500 mM NaCl, 50 mM Tris-HCl at pH 8.0, 50 mM EDTA, and 4% sodium dodecyl sulfate) followed by cleaning using QIAamp Fast DNA Stool Mini Kit DNA and stored at −20°C until processing. All four samples were sequenced using an Illumina HiSeq 2500 sequencer with a read length of 250 bp.

### Bacteriome data

Bacteriome data used in this study were obtained from a previous investigation [11]. In this study, faecal samples from 80 WHS mice were collected and analysed using 150 bp paired-end reads generated by shotgun metagenome sequencing. This approach enabled the generation of 374 high-quality bacterial metagenome-assembled genomes (MAGs) by employing various sequence assembly and binning methods. The relative abundance of each bacterial MAG in each sample was calculated by mapping quality-filtered sequences to the MAG database using CoverM v0.6.1 (https://github.com/wwood/CoverM). The relative abundance data from the earlier study were subsequently reused for analysis in the current study.

### Quality control of metagenomic sequences

All sequences obtained were quality-filtered using KneadData (https://huttenhower.sph.harvard.edu/kneaddata/), where low-quality sequences and any contamination from PhIX, host (mouse_C57BL_6NJ.1), human genome (hg37dec_v0.1.1), and transcriptome (human_hg38_refMrna.1) were discarded. Kneaddata uses Trimmomatic[16] to remove adapters, TRF[17] to remove repetitive sequences, and bowtie2[18] to remove contamination.

### Viral sequences generation and predication

In this study, we utilized quality-filtered shotgun metagenome data from both newly generated (4 samples; 250 bp reads) and previously published datasets (80 samples; 150 bp reads)[11]. Unless otherwise stated, all analyses were performed using default settings. Filtered sequences were assembled using metaSPAdes v3.15.3 (-k 21,33,55,77,99,127)[19] in the single-sample assembly mode. The assembled sequences were then analysed with Phamb v1.0.1[20], geNomad v1.8.0 (end-to-end --enable-score-calibration --composition metagenome)[21], Phables v1.3.2[22], and VirSorter2 v2.2.4 (--include-groups "dsDNAphage,NCLDV, RNA,ssDNA,lavidaviridae" --keep-original-seq)[23] for independent viral sequence predications. The geNomad output was further processed using COBRA v1.2.3[24] and directly fed into CheckV v1.0.3(checkv-db-v1.5)[25] for quality assessment. Simultaneously, metaViralSpades v3.15.3[26] was used for the single-sample assembly of viral sequences from all samples, and MetaHipMer2 v2.2.0.0.3[27] was employed to co-assemble 250 bp reads from four samples. The output from MetaHipMer2 was analysed using GeNomad for viral sequence prediction. In total, seven independent inputs were fed into CheckV for quality checking: VirSorter2, GeNomad, Phamb, Phables, COBRA (processed from geNomad), MetaViralSpades, and the MetaHipMer2-GeNomad pipeline.

We followed previous protocols for quality filtering[28, 29]. Briefly, CheckV was run in “end_to_end” mode with default settings on all viral sequences. We only kept those sequences which met the following criteria (1) with higher number of viral genes than host genes, (2) longer than 3 kbp, (3) the viral sequences are represented in the contig less than/equal to one (kmer_freq□≤□1), (4) without warning “>1 viral region detected” and “contig >1.5× longer than expected genome length”, and (5) completeness is more than 25%. Overall, 60588 viral sequences met these criteria. Additionally, sequences which remained “undetermined” but longer than 3kbp were combined and fed into geNomad v1.8.0 (end-to-end --enable-score-calibration --sensitivity 7 --composition metagenome) with stringent settings. We obtained 1428 sequences predicted to be viral using geNomad.

Further we removed all sequences which contains prokaryotic housekeeping marker genes using fetchMG (v1.0)[30], and ribosomal RNA genes of bacteria, archaea, eukaryota, and metazoan mitochondria using barrnap (0.9) (https://github.com/tseemann/barrnap).

### Clustering of viral sequences into viral operational taxonomic units (vOTUs)

The nucleotide BLAST database of all viral sequences was built by makeblastdb (option “-dbtype nucl”) from Blast v2.12.0, and the pairwise comparisons were generated by blasting all viral sequences all-against-all with blastn (option “-max_target_seqs 10,000”). Afterwards, two custom scripts (anicalc.py and aniclust.py) from the CheckV repository (https://bitbucket.org/berkeleylab/checkv/src/master/) were used to compute ANI and AF for clustering into species-level viral operational taxonomic units (vOTUs) on the basis of 95% ANI and 85% AF of the shorter sequence (options “-min_ani 95, - min_tcov 85, -min_qcov 0”)[31]. The longest viral sequences were selected as cluster representatives of the vOTUs.

### Taxonomic assignment and functional annotation of vOTUs

Taxonomic assignment was done using geNomad v1.8.0 with the “annotate” command with MMseqs2[32] sensitivity set at “10”. For functional annotation, prodigal-gv v2.22.0[21, 33] available at (https://github.com/apcamargo/prodigal-gv) with “-p meta” command was used.

### Viral host prediction

For viral host prediction, we used iPHoP v1.3.3[34] which combines the output from blastn[35] with the host genomes, blastn[35] with the CRISPR spacer database, Random Forest Assignment of Hosts (RaFAH)[36], WIsH[37], VirHostMatcher (VMH)[38, 39], and prokaryotic virus Host Predictor (PHP)[40]. The standard iPHoP database (iPHoP_db_Aug23_rw) was used for the predictions. All predication with “Confidence score” of more than “90.00” are considered to be true host predications.

### Determining relative abundance of vOTUs representatives in metagenomes

To determine the relative abundance of viral sequences in faecal metagenomic samples, we used reads per kilobase per million mapped reads (RPKM) values estimated using CoverM v0.7.0. We built a mapping database using the WHS-MV catalogue in BWA v2.2.1[41]. Quality-controlled reads from each sample were mapped using CoverM with the options set to ensure high-quality mapping (--min-read-aligned-percent 95, --min-read-percent-identity 90, --min-covered-fraction 75, -- m rpkm). Reads with less than 90% identity were excluded and the minimum coverage threshold for vOTU presence was set at 75%. The RPKM values obtained for vOTU representatives within each sample were normalised to the total RPKM value to calculate percentages. In addition, the mapping rate of quality-controlled reads for each sample was obtained from the CoverM output.

### Statistical analysis

RStudio (https://www.rstudio.com/) was used for all the statistical analysis with R-4.4.1 (https://cran.r-project.org/). The details of the statistical tests are provided in each figure legend.

#### Diversity correlation analysis

To analyse the relationship between bacteriome and virome diversity, both regression and Pearson’s correlation analyses were performed. The data were visualised using various R packages, including Corrplot v09.2[42] and ggpubr v0.6.0[43].

#### PERMONOVA analysis

Microbial community data were analysed using the vegan package v2.6-6.1 in R v4.4.1[44]. The environmental variables analysed included bacterial Shannon diversity, bacteriome Bray-Curtis distance, active contact time, sex, mouse group (S1, S2, C1, C2), and selection (tameness group or control group). The categorical variables (sex, mouse group, and selection) were converted into factors. Bray-Curtis dissimilarity was used to calculate the distance matrix. PERMANOVA was performed using adonis2 function.

#### Mantel R analysis

Microbial community data were analysed using the vegan package v2.6-6.1 in R v4.4.1[44]. The environmental variables analysed included bacterial Shannon diversity, bacteriome Bray-Curtis distance, active contact time, sex, mouse group (S1, S2, C1, C2), and selection (tameness group or control group). Abundant data were extracted, and categorical variables (sex, mouse group, and selection) were converted to numeric values. Bray-Curtis dissimilarity was used to create a distance matrix from the abundance data. Mantel tests were performed to assess the correlations between this matrix and the Euclidean distance matrices of each environmental variable, using 9999 permutations for significance testing. P-values were adjusted for false discovery rate (FDR).

#### vOTUs and bacteria abundance correlation analysis

Bacterial abundance and taxonomic data were loaded and merged. Virome abundance and virome-host prediction data were similarly loaded and merged. Merged bacterial data were aggregated at the genus level. Virome data were grouped according to the predicted host genus. Both datasets were filtered to include relevant genera and sorted by genus[45–47]. Spearman correlations between virome and bacterial abundance were computed for each genus using the Hmisc v5.1-3 package[48]. The Benjamini-Hochberg method was applied for false discovery rate (FDR) adjustment using ‘multtest’ v2.60 package[49].

## Results

### Gut virome database of WHS mice

To comprehensively analyse the gut virome in WHS mice, we used 80 publicly available shotgun metagenome datasets from our previous study[11]. Additionally, we generated four extra samples of 250 bp paired-end sequences in the current study. Given the complexity and low abundance of the gut virome, we employed multiple independent methods to identify the viral sequences (Supplementary Fig. 1). In total, 1,405,278 viral sequences were obtained. After stringent quality filtering using CheckV, followed by rRNA sequence and bacterial marker gene removal, 59,853 viral sequences were retained. We decided to set the cutoff at 25% completeness in CheckV, rather than 50%, to incorporate as many unique viral sequences as possible into our database. Clustering of these viral sequences with 95% sequence similarity generated 6094 viral vOTUs[31] (corresponding to the species level).

Subsequently, we removed 15 vOTUs assigned to the viral families *Phycodnaviridae* (n = 5), Marseilleviridae (n = 5), *Mimiviridae* (n = 4), and *Iridoviridae* (n = 1) due to their likely representation of contaminants or misclassifications, as previously suggested. One vOTU from the *Nanoviridae* family was also removed because its natural hosts are plants that likely originate from mouse feed. The remaining 6,078 vOTUs were designated as the WHS mouse virome (WHS-MV) catalogue (Fig. 1A). Among these, 25 vOTUs were categorised as complete, 368 vOTUs as high quality (with more than 90% completeness, including 100 vOTUs with 100% completeness), 699 vOTUs as medium quality (50–90% completeness), 1,294 vOTUs as low quality (25–49.99% completeness), and 3,692 vOTUs remained as “Not-determined” (Fig. 1B) from checkV analysis.

**Fig. 1.**
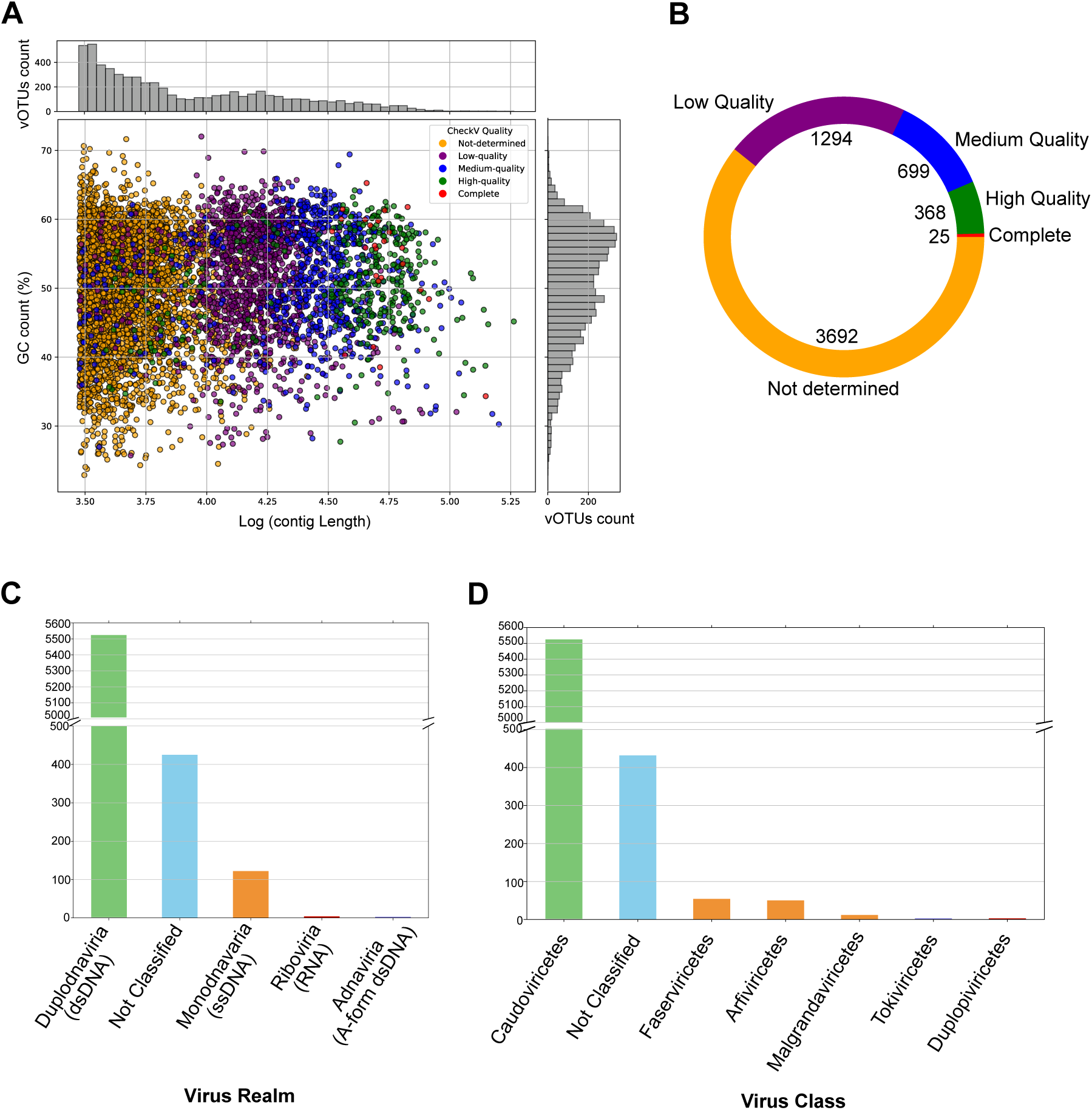
Overview of 6,078 vOTUs constructed in the present study. **(A)** Genome size and GC content of vOTUs reconstructed from the dataset of 84 shotgun metagenome samples. Bar plots on the top and right side depict the distribution of genome size and GC content, respectively. Each vOTUs is coloured based on CheckV quality results. **(B)** CheckV quality results of all vOTUS. **(C)** Taxonomy result count of all vOTUs at realm level. **(D)** Taxonomy result count of all vOTUs at class level.

GeNomad, which provides reliable taxonomic information, was used for taxonomic annotation. Of the 6,078 vOTUs, 5,660 were annotated as viruses, with 5,653 having taxonomic information at the realm level; 5,640 at the kingdom, phylum, and class levels; 128 at the order level; and 178 at the family level. Taxonomically, 5,524 vOTUs belonged to the *Duplodnaviria* (dsDNA) realm, 122 to the *Monodnaviria* (ssDNA) realm, 4 to the *Riboviria* (RNA-dependent) realm, and 3 to the *Adnaviria* (A-form dsDNA) realm (Fig. 1C). Taxonomic information was not obtained for 418 vOTUs. All 5,524 *Duplodnaviria* vOTUs belonged to the *Caudoviricetes* class, a well-known bacteriophage class. In *Monodnaviria*, 54 vOTUs belonged to the *Inoviridae* family *(Faserviricetes*), 12 to the *Microviridae* family (*Malgrandaviricetes*), and 50 to the *Circoviridae* family (*Arfiviricetes*). Only the *Inoviridae* and *Microviridae* families are known bacteriophages. In *Riboviria*, three vOTUs belonged to the *Partitiviridae* family (*Duplopiviricetes*). All three *Adnaviria* vOTUs belong to the *Lipothrixviridae* family (*Tokiviricetes*), which is known to infect hyperthermophilic archaea[50] (Fig. 1D). Additionally, we identified one vOTU belonging to the *Bicaudaviridae* family, which infects hyperthermophilic archaea[51].

In summary, we created a WHS-MV catalogue that includes various viruses and bacteriophages with diverse genetic materials, such as dsDNA, ssDNA, RNA, and A-form dsDNA (Supplementary Table 1). This catalogue will be a valuable resource for future studies on the gut virome of wild-derived mice, aiding the discovery of new viruses.

### Phage host analysis

Before performing phage host prediction, we filtered out vOTUs from the *Circoviridae* family because animals are their natural hosts[52]. Consequently, host predictions were conducted for 6028 vOTUs, resulting in successful predictions for 3307 phages. Among these, 3065 phages were found to infect only one host genus (specialist phages), whereas 242 phages were identified as generalist phages capable of infecting more than one bacterial genus. Of these generalists, 66 could infect more than one family of bacteria, 11 could infect more than one order, and 2 could infect two different phyla (Fig. 2).

**Fig. 2.**
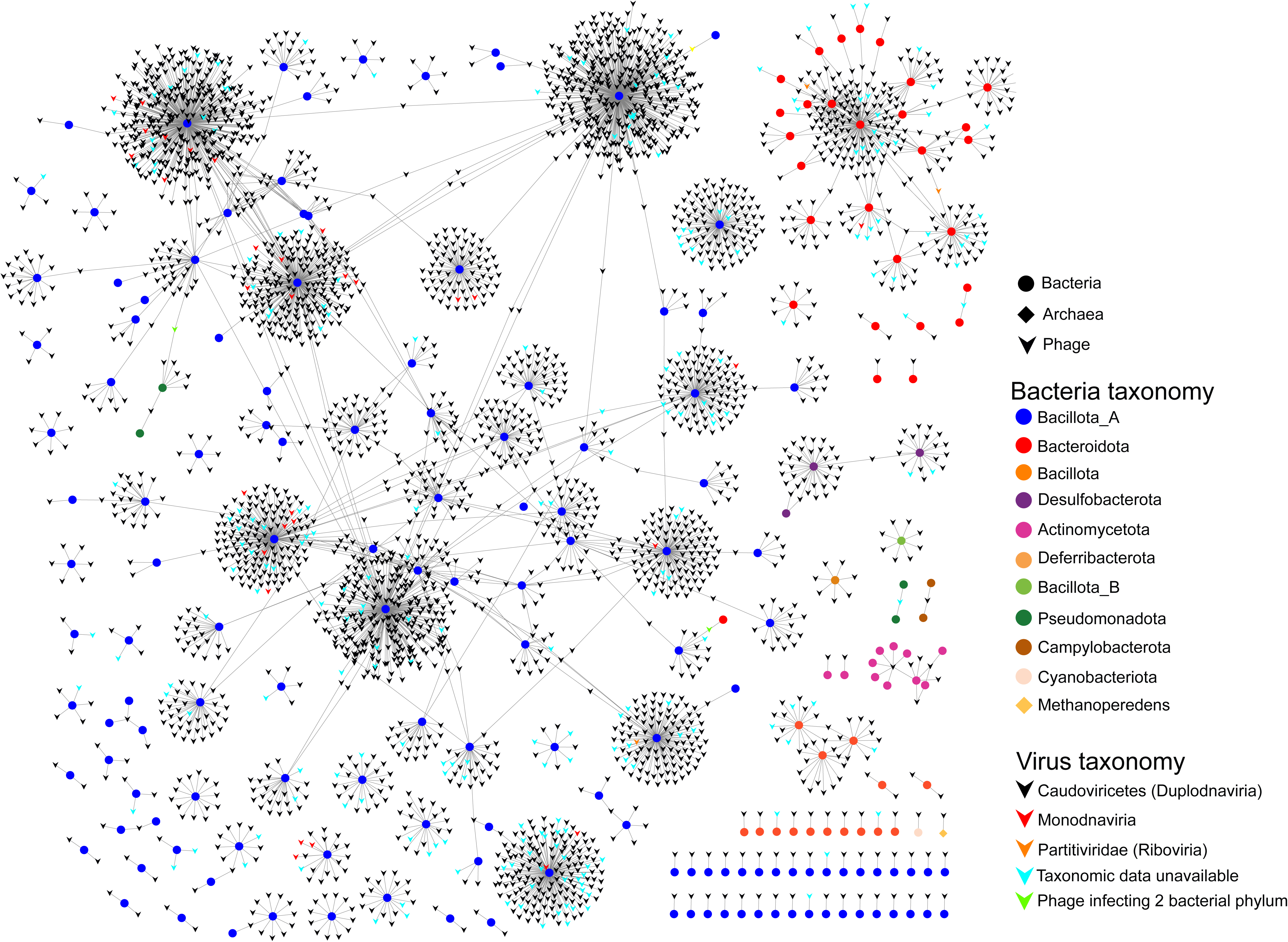
Network graph of all phage-host interaction in this study.

In terms of host taxonomy, 2917 phages infect *Bacillota_A*, 282 infected *Bacteroidota*, 52 infected *Bacillota*, 51 infected *Desulfobacterota*, and eight each infected *Actinomycetota, Bacillota_B*, and *Pseudomonadota*. Notably, one phage infected *Campylobacterota, Cyanobacteriota*, and *Halobacteriota*. *Halobacteriota*, an archaeon, are infected by phages belonging to the *Caudoviricetes* family (Table 1).

**Table 1.** Phage host taxonomy and the count of phages infecting each bacterial phylum.

| Phylum | Number of vOTUs | Generalist | Specialist |
| --- | --- | --- | --- |
| *Actinomycetota* | 8 | 3 | 5 |
| *Bacillota* | 51 | 9 | 42 |
| *Bacillota_A* | 2891 | 139 | 2752 |
| *Bacillota_B* | 8 | 0 | 8 |
| *Bacteroidota* | 282 | 86 | 196 |
| *Campylobacterota* | 1 | 1 | 0 |
| *Cyanobacteriota* | 1 | 0 | 1 |
| *Deferribacterota* | 7 | 0 | 7 |
| *Desulfobacterota* | 51 | 3 | 48 |
| *Halobacteriota* | 1 | 0 | 1 |
| *Pseudomonadota* | 8 | 3 | 5 |

Additionally, we discovered two phages from the *Caudoviricetes* order capable of infecting two phyla: one infecting *Bacillota_A* and *Bacteroidota*, and the other infecting *Bacillota_A* and *Pseudomonadota* (Fig. 2). There are already few reports available on phages capable of infecting multiple phyla [53, 54].

Regarding the taxonomy of vOTUs, 3031 belonged to *Duplodnaviria*, 34 belonged to *Monodnaviria*, and three were RNA viruses (*Riboviria*). We lacked taxonomic information on 239 vOTUs. All 3031 vOTUs in *Duplodnaviria* belonged to the *Caudoviricetes* order. Of the 34 vOTUs in *Monodnaviria*, 30 were identified as belonging to the family *Inoviridae*, all of which infect *Bacillota_A*, while 1 vOTU belonged to *Microviridae*, infecting *Bacteroidota*. All three *Riboviria* vOTUs belonged to the *Partitiviridae* family, with two infecting *Bacteroidota* and one infecting *Bacillota_A* (Table 2).

**Table 2.** Taxonomy and distribution of phages.

| Realm | Number of vOTUs | Generalist | Specialist |
| --- | --- | --- | --- |
| Duplodnaviria | 3,031 | 236 | 2,795 |
| Monodnaviria | 34 | 0 | 34 |
| Riboviria | 3 | 1 | 2 |
| No Taxonomy Information | 239 | 5 | 234 |

This analysis provides a comprehensive overview of phage-host interactions in the WHS mouse gut virome, providing valuable insights for future research.

### Tameness selection pressure leads to changes in gut virome beta diversity

To investigate the effect of selective breeding for tameness on the gut virome profile of WHS mice, we analysed 80 samples of 150bp reads. This focus was chosen because previous analyses of the bacterial influence on tameness were conducted using these samples[11]. Abundance was calculated using CoverM, revealing that, on average, 5% of the reads mapped to the WHS-MV catalogue, with a range from 3.31% to 11.69% (Supplementary Fig. 2). Relative abundance, measured as RPKM values, was used for diversity analysis.

Our analysis showed a similar virome profile across all four groups at the class and realm levels, with *Caudoviricetes* (*Duplodnaviria*) accounting for over 80% of the viral abundance in the WHS mouse gut (Fig. 3 A, B). There were no significant differences in α-diversity (Shannon diversity) among the groups (Fig. 3C), although the C1 group exhibited slightly higher Shannon diversity compared to the others. Interestingly, both selected groups demonstrated higher β-diversity (Bray-Curtis distance) compared to the control groups (Fig. 3D, E; Supplementary Table 2), suggesting that tameness selection pressure leads to greater dissimilarity in the gut virome of the selected mice groups relative to the controls.

**Fig. 3.**
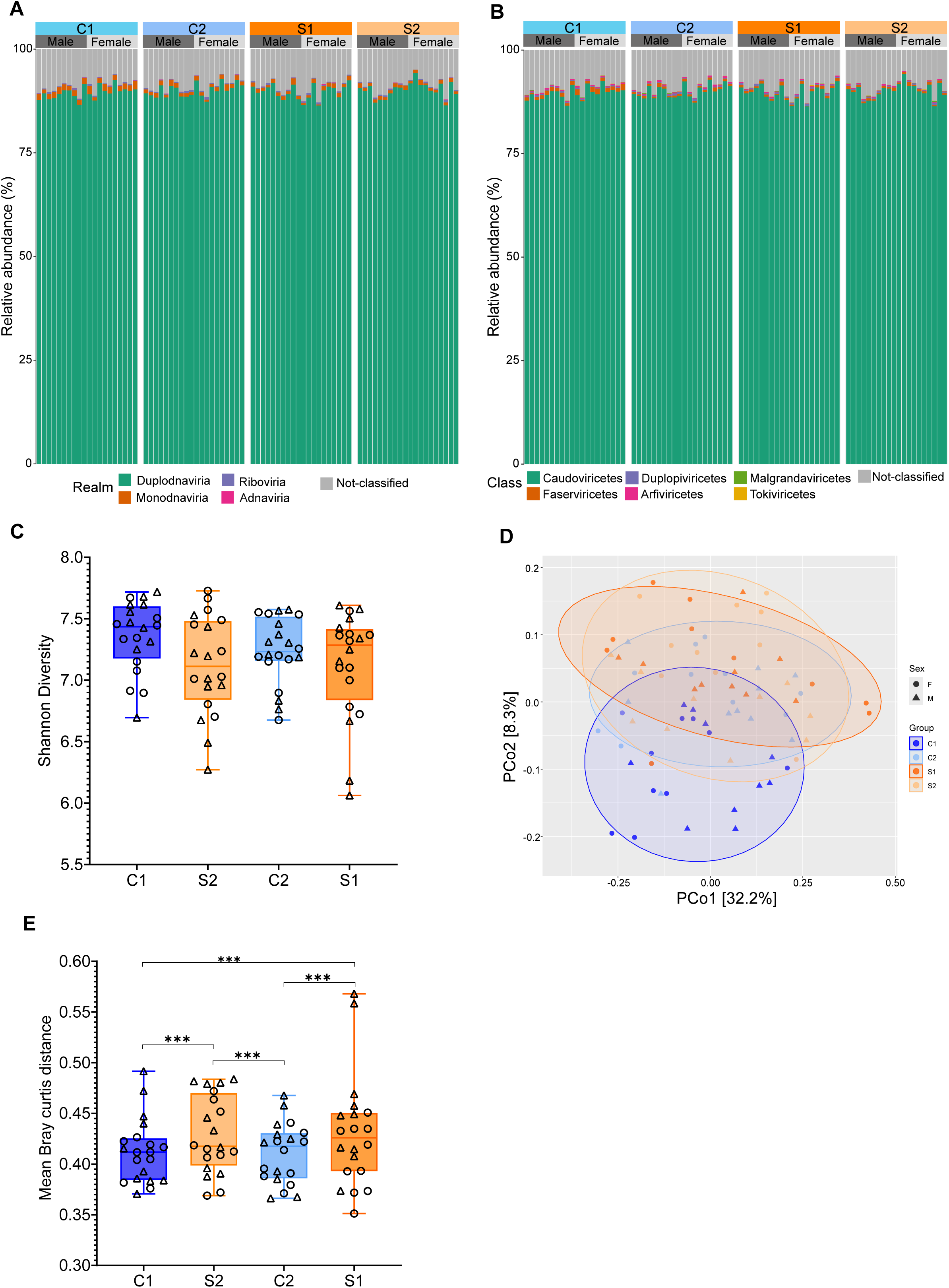
Gut virome diversity in WHS mice groups. **(A)** Realm level relative abundance across all groups. **(B)** Class level relative abundance across all groups. **(C)** Shannon diversity index across four groups of WHS mice. **(D)** Beta diversity based on Bray– Curtis dissimilarity. **(E)** Box plot of mean bray–curtis distance in all four groups. Asterisks represent statistical significance (***p<0.001, Wilcoxon rank-sum test, two-sided). N= 80 samples (20 in each group with 10 male and 10 female).

We further compared α-diversity (Shannon diversity) and β-diversity (Bray-Curtis distance) between bacteriome and the virome. For bacterial data, we used previously generated WHS metagenome-assembled genomes[55]. Notably, species-level Shannon diversity (Fig. 4A, R=0.75, p=8.4e-16) and mean Bray–Curtis distances (Fig. 4B, R=0.71, p=2.1e-13) showed a significant Pearson correlation between the bacteriome and virome, indicating a close relationship between the virome and bacteriome structures in the gut. Additionally, the β-diversity of the virome was significantly higher than that of the bacteriome (Fig. 4C, p<0.001), suggesting that the virome is more specific to each individual than the bacteriome.

**Fig. 4.**
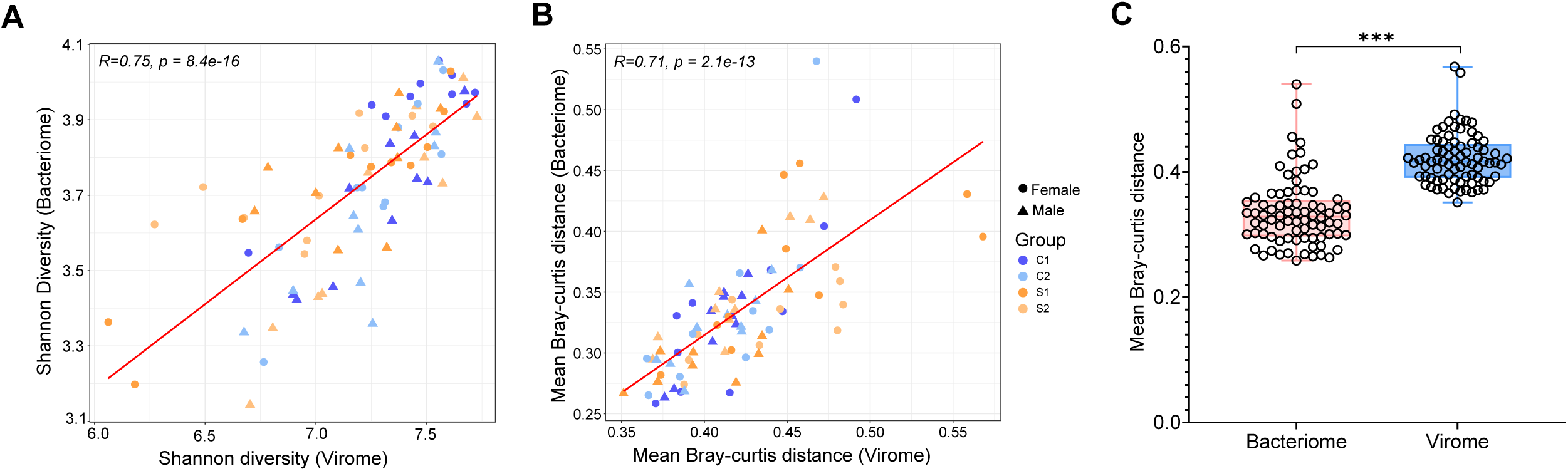
Correlation between bacteriome and virome diversity **(A)** α-diversity (Shannon index) Pearson correlation between bacteriome and virome. **(B)** Mean β-diversity (Bray-Curtis distance) Pearson correlation between bacteriome and virome. **(C)** Box plot of mean bray-curtis distance between bacteriome and virome. Asterisks represent statistical significance (***p<0.001, Wilcoxon rank-sum test, two-sided). N= 80 samples (20 in each group with 10 male and 10 female).

### Phage vOTUs abundance correlation with their host abundance

To further explore the association between phages and their hosts, we examined one-to-one correlations between the relative abundance of each phage and its predicted host at the genus level. We found a positive correlation, with an average Spearman correlation of 0.397, suggesting that phages and host bacterial species co-occur, rather than being mutually exclusive, in the WHS mouse gut. Focusing on phages that infect *Limosilactobacillus* and *Gallimonas*, based on our previous findings[11], we identified 17 phages that infect *Limosilactobacillus* and 21 phages which infect *Gallimonas* genus. Among the phages infecting *Limosilactobacillus*, 5 were generalists and 12 were specialists. There were 14 significant positive correlations between the phages and *Limosilactobacillus* abundance (Fig. 5A). All phages infecting *Gallimonas* were specialists and exclusively infected *Gallimonas*, with 14 phages showing significant positive correlations (Fig. 5B). To further assess the overall influence of these phages on the abundance of *Limosilactobacillus* and *Gallimonas*, we combined the abundance data for all phages that infected these bacterial genera. We observed a significantly high positive correlation for both *Limosilactobacillus* (R=0.66, p=4.8E-11) and *Gallimonas* (R=0.85, p= 5.9E-23) (Figure 5). This implies that the populations of both *Limosilactobacillus* and *Gallimonas* are controlled by phages, which might indirectly influence tameness behaviour in WHS mice.

**Fig. 5.**
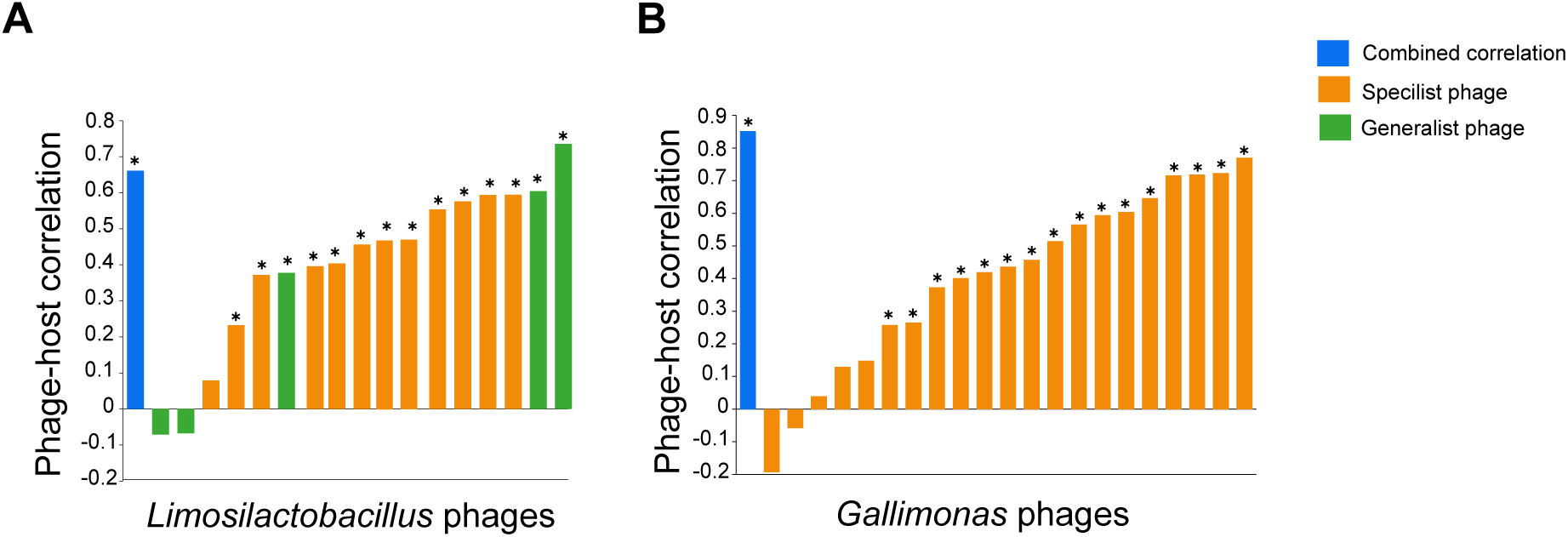
Correlation between bacterial MAGs and viral OTUs abundance. **(A)** Spearman correlations between the relative abundances of *Limosilactobacillus* infecting phages and *Limosilactobacillus* genus. **(B)** Spearman correlations between the relative abundances of *Gallimonas* infecting phages and *Gallimonas* genus. Asterisks (*) denote statistical significance (FDR□<□0.05). N = 80 samples (20 in each group, with 10 males and 10 females).

### Factors shaping WHS mice gut virome

To determine the extent to which different factors affect the WHS mouse gut virome, we selected six metadata: bacteriome Shannon diversity, bacteriome Bray-Curtis distance, active contact time, gender, mouse group (S1, S2, C1, C2), and selection (tame group or control group). Thus, we first performed both PERMANOVA and MANTEL analyses based on the virome Bray–Curtis distances to estimate the effect size and correlation coefficient of all metadata factors. Both analyses revealed a significant (FDR□<□0.05, 1000 permutations) contribution and correlation between the WHS mouse gut virome and all metadata factors, except active contact time (Fig. 6A for PERMANOVA analysis and Fig. 6B for MANTEL analysis). Shannon diversity accounted for the highest viral taxonomic variation (22.33%; Fig. 6A). Additionally, we found that the mouse group (S1, S2, C1, C2), bacteriome Bray-Curtis distance, selection (tameness group or control group), and sex had systematic effects on the overall composition of the viral community, with 7.08%, 4.98%, 4.40%, and 2.18% of variance, respectively. A similar pattern was found in the MANTEL analysis, with all metadata, except active contact time, showing a significant correlation with virome Bray–Curtis distances (Fig. 6B).

**Fig. 6.**
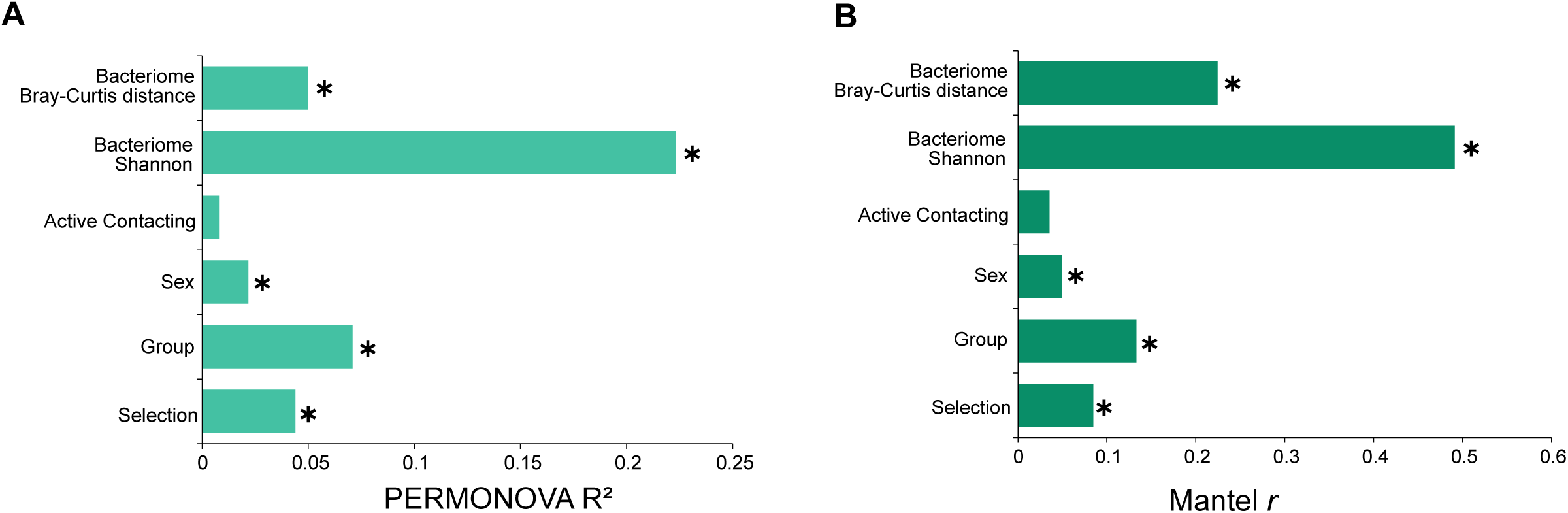
Factors shaping the WHS mice gut virome during selective breeding. **(A)** Effect size (R^2^) of each metadata as determined by PERMANOVA. **(B)** Mantel r correlation of each metadata with the mean bray-curtis distance of the virome. Asterisks (*) denote statistical significance (FDR□<□0.05) of each factor. N = 80 samples (20 in each group, with 10 males and 10 females).

## Discussion

Our study provides a comprehensive analysis of the gut virome in wild-derived heterogeneous stock (WHS) mice selectively bred for tameness. These findings highlight the significant impact of selective breeding on the gut virome, demonstrating that alterations in microbial communities extend beyond bacteria to viruses. This underscores the intricate relationship between the gut microbiome, including its viral components, animal host behaviour, and physiology.

One of the key findings of this study was the difference in gut virome beta diversity between the selectively bred and control groups. The higher beta diversity in the tameness-selected groups suggests that selective breeding for tameness exerts distinct pressure on the gut virome, leading to a more heterogeneous viral community. This supports the hypothesis that selective breeding for behavioural traits can influence the composition and diversity of the gut virome, potentially through changes in the bacterial community that serves as a host for these viruses[11, 56, 57].

Our analysis of phage-bacterial host interactions revealed a strong correlation between the abundance of specific phages and their bacterial hosts. This indicates co-occurrence rather than mutual exclusivity of phages and their bacterial hosts in the WHS mouse gut, suggesting that the gut virome is closely tied to the bacterial community structure[58, 59]. The significant positive correlations between the abundance of phages infecting *Limosilactobacillus* and *Gallimonas* and their respective bacterial hosts further highlight the potential role of phages in regulating bacterial populations and consequently influencing host behaviour. This aligns with our previous findings, in which increased levels of *Limosilactobacillus reuteri* were associated with enhanced tameness and elevated blood oxytocin levels[11].

The correlation between Shannon diversity and Bray-Curtis distances of the virome and bacteriome suggests a close interrelationship between these microbial communities[60, 61]. The higher beta diversity of the virome compared to that of the bacteriome indicates that the viral component of the gut microbiome is more individualised, potentially reflecting the unique viral-bacterial interactions within each bacterial host. This specificity could be a result of selective pressures exerted by both the immune system of the animal host [62] and bacterial community dynamics, which are influenced by selective breeding for tameness.

The identification of diverse vOTUs and their taxonomic classifications provide a valuable resource for understanding the role of the gut virome in domestication processes. The presence of phages that can infect multiple bacterial genera and even different bacterial families and orders suggests that these phages may play a role in modulating the structure and function of the gut microbiome. This modulation can influence the physiological and behavioural traits of the host, contributing to the domestication process. The significant contribution of bacterial diversity and composition to the taxonomic variation in the gut virome further supports the notion that the microbial ecosystem, including both bacteria and viruses, is integral to the domestication process.

Although this study provides valuable insights into the role of the gut virome in selective breeding and domestication of host animals, there are limitations that warrant further investigation. Reliance on shotgun metagenome sequencing and bioinformatics predictions may not capture the full complexity of viral-bacterial interactions and their functional implications. Future studies should incorporate experimental validations of phage-bacterial host interactions and explore the functional roles of the identified phages in modulating the behaviour and physiology of the host animal. In addition, expanding the analysis to other selectively bred animal models and exploring the longitudinal effects of selective breeding on the gut virome could provide a more comprehensive understanding of the mechanisms underlying microbial contributions to domestication. Investigating the effect of the gut virome on other aspects of the host animal’s health, such as immunity and metabolism, could also reveal the broader implications of the gut microbiome in shaping animal traits.

This study advances our understanding of the role of the gut virome in selective breeding and domestication by revealing the impact of tameness selection on viral diversity and phage-bacterial host dynamics. These findings highlight the intricate interplay between the gut virome and bacteriome, suggesting that phages are integral components of the gut microbiome that contribute to the modulation of host animal behaviour and physiology. This study lays the foundation for future studies aimed at unravelling the complex microbial interactions that influence animal domestication and selective breeding outcomes.

## Acknowledgments

We thank Motoko Nihei and Akiko Tsuchiya for their technical assistance in this study. Computations were partially performed on the NIG supercomputer at ROIS National Institute of Genetics.

## Author contributions

B.B.B. and T.K. conceived the project; B.B.B. and T.K. designed the study; B.B.B. and K.F. performed the analysis; A.T. performed shotgun metagenome sequencing; T.K. provided supervision and funding; B.B.B. and T.K. wrote the manuscript; all authors provided comments and feedback on the manuscript. All the authors approved the final version of the manuscript.

## Conflicts of interest

Authors has no competing interests regarding this research.

## Funding

This research was supported by a grant from JST SPRING (Grant Number JPMJSP2104) awarded to B. B. and a Grant-in-Aid for Scientific Research (JSPS KAKENHI Grant Numbers 19KK0177 and 24K01951) awarded to T.K.

## Data and code availability

All data required to evaluate the conclusions in the paper are presented in the paper and/or supplementary data. Raw shotgun metagenomic sequencing datasets generated in this study are available at the NCBI under BioProject PRJDB18588, with BioSample accession numbers SAMD00805331 to SAMD00805334 and SRA accession numbers DRR585793 to DRR585796. The non-redundant vOTUs generated in this study are available in the Zenodo database (https://doi.org/10.5281/zenodo.13220406). No new code was generated in this study.

## Notes

https://doi.org/10.5281/zenodo.13220406

### Competing Interest Statement

The authors have declared no competing interest.

