## Supplementary Figures for "Tameness selection pressure affects gut virome diversity in mice"

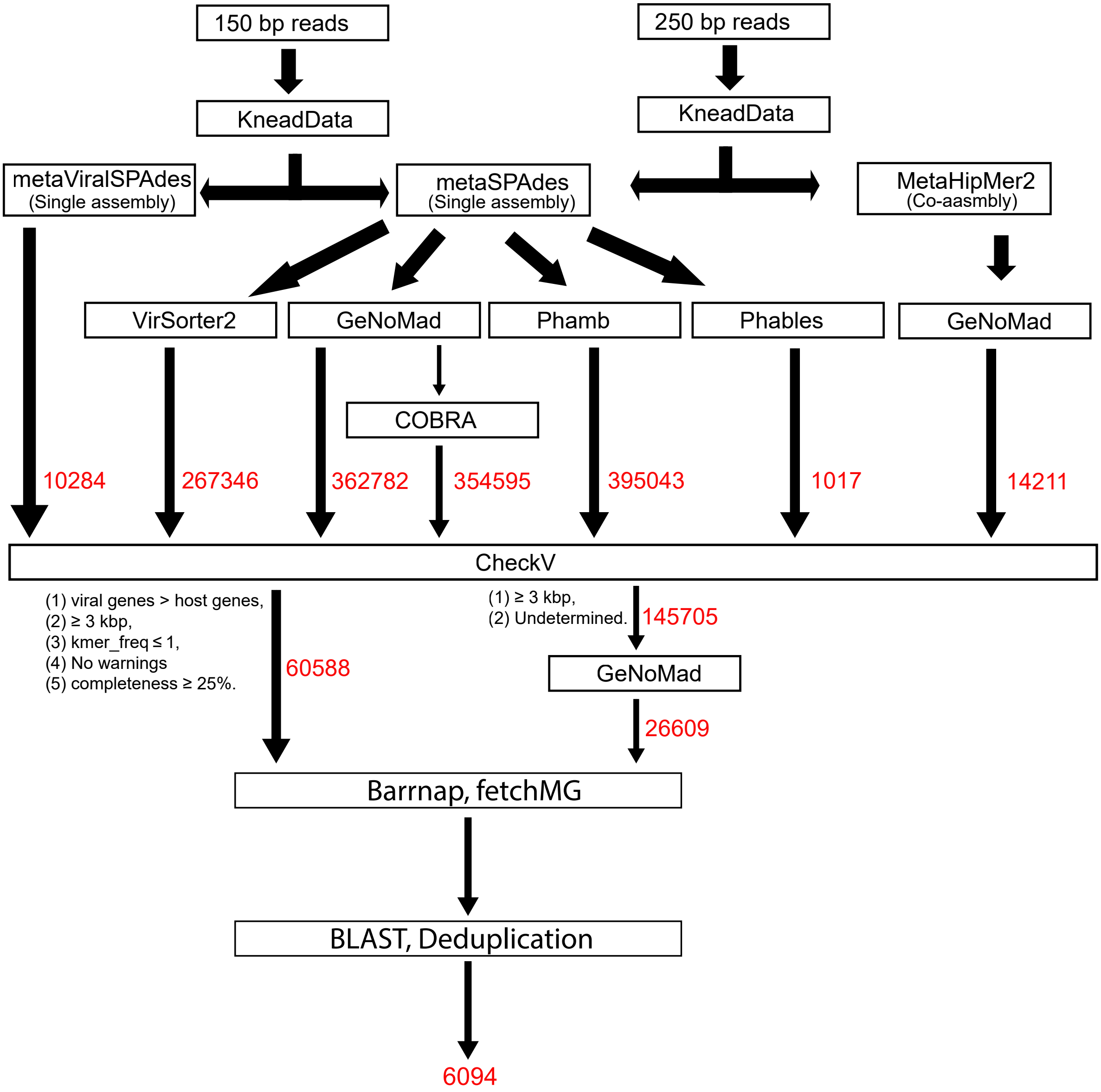


Supplementary Fig. 1: Schematic of vOTU Generation from Shotgun Metagenome Data. Sequences generated at each step are highlighted in red.

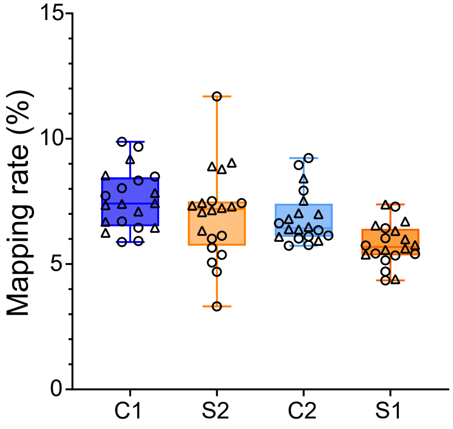


Supplimentary Fig. 2. Mappping rate of quality filtred reads from each sample against WHS-MV catalogue.
N = 80 (20 per group (10 male and 10 female))
